# Associative coding of conditioned fear in the thalamic nucleus reuniens in rodents and humans

**DOI:** 10.1101/2025.03.18.643915

**Authors:** Tuğçe Tuna, Michael S. Totty, Muhammad Badarnee, Flávio Afonso Gonçalves Mourão, Shaun Peters, Mohammed R. Milad, Stephen Maren

## Abstract

The nucleus reuniens (RE) is a midline thalamic structure interconnecting the medial prefrontal cortex (mPFC) and the hippocampus (HPC). Recent work in both rodents and humans implicates the RE in the adaptive regulation of emotional memories, including the suppression of learned fear. However, the neural correlates of aversive learning in the RE of rodents and humans remains unclear. To address this, we recorded RE activity in humans (BOLD fMRI) and rats (fiber photometry) during Pavlovian fear conditioning and extinction. In both rats and humans, we found that conditioned stimulus (CS)-evoked activity in RE reflects the associative value of the CS. In rats, we additionally found that spontaneous neural activity in RE tracks defensive freezing and shows anticipatory increases in calcium activity that precede the termination of freezing behavior. Single-unit recordings in rats confirmed that individual RE neurons index both the associative value of the CS and defensive behavior transitions. Moreover, distinct neuronal ensembles in the RE encode fear versus extinction memories. These findings suggest a conserved role of the RE across species in modulating defensive states and emotional memory processes, providing a foundation for future translational research on fear-related disorders.

## INTRODUCTION

Decades of work reveal a critical role for the medial prefrontal cortex (mPFC) and the hippocampus (HPC) in the regulation of emotional memory, including fear conditioning, extinction, and extinction retrieval processes on both rodents and humans^1–10^. This work supports a model by which contextual information encoded by the HPC orchestrates mPFC-dependent memory retrieval functions^11–13^ via direct projections from the HPC to the mPFC^14–17^. Interestingly, the mPFC also influences memory retrieval functions by the HPC^18–20^, though the neural circuit mechanism for this is unclear. The thalamic nucleus reuniens (RE), a ventral midline structure located above the third ventricle that interconnects the mPFC and HPC^21–23^, may mediate this function^24–27^.

Recent work in both rodents and humans implicates the RE in the adaptive regulation of emotional memories, including the suppression of conditioned fear^24,28–31^. In rodents, both the RE and its projections with the HPC are critical for both the acquisition and extinction of contextual fear memories^32–35^. Moreover, selective silencing of mPFC➙RE projections attenuates extinction learning and retrieval^30^, and optogenetic modulation of the RE is necessary and sufficient for both the retrieval of extinction memories and mPFC-HPC theta synchrony^36^. Although these results demonstrate that RE is critical for the regulation of fear and extinction memories in rodents, the function of the RE in humans remains largely unexplored due to challenges imaging midline thalamic nuclei in the human brain. In addition, neuronal activity in the RE during different phases of fear learning has not been characterized in either rats or humans.

To address these issues, we used fear conditioning, extinction, and extinction retrieval procedures in healthy humans and wild-type rats to examine the functional homology of the RE across species. Using blood oxygenation level dependent (BOLD) functional magnetic resonance imaging (fMRI) in humans and bulk fiber photometry in rodents, we found that the RE activity in rats and humans encodes the associative value of the CS across fear conditioning, extinction, and extinction retrieval. In rodents, we also found that RE activity closely tracks defensive states and anticipates reductions in freezing behavior. Using *in vivo* electrophysiology with unsupervised clustering methods, single-unit recordings in rats confirmed that individual RE neurons track both the associative salience of the CS and defensive state transitions. In addition, we found that distinct RE ensembles encode fear versus extinction memories. Together, the RE shows divergent, and possibly independent, neural signatures during fear conditioning: one encoding stimulus salience and the other encoding defensive states. These findings suggest a conserved role of the RE across species in encoding fear memory processes, providing a foundation for future translational research on fear-related disorders.

## RESULTS

### BOLD activity in the human RE encodes associative value during fear conditioning and extinction

To determine the contribution of RE to fear learning in humans, humans underwent a discriminative fear conditioning in distinct computer-generated visual contexts as previously described^10,37^. We analyzed the early stage of learning (the first 4 trials), when the CS-US association is acquired in humans. We first compared the averaged BOLD signal to CS+ vs. CS-across all 4 trials. We then performed a trial-by-trial comparison of BOLD signal to CS+ and CS-, which enables a higher temporal resolution characterization of the activation patterns. Here, we only report BOLD signal from the RE. Behavioral results can be found in a previous publication^38^.

During conditioning, we observed a differential level of BOLD activity to the CS+ and CS-within the RE [*t*_292_ = 5.53, *p* < 0.001, mean difference 95 % CI (0.12 - 0.24), Cohen’s *d* = 0.32 with 95 % CI (0.21 - 0.44)]. We also found a consistent increased signal across trials, with a heightened response to CS+ during the second trial (main effect of Trial: *F*_2.2,656.8_ = 3.45, *p* = 0.027, partial *η*² = 0.012; main effect of CS: *F*_1,292_ = 30.62, *p* < 0.001, partial *η*² = 0.095; no Trial x CS interaction: *F*_2.6,783.1_ = 1.38, *p* = 0.25, partial *η*² = 0.005. Post-hoc_FDR corrected_ for CS+ > CS-: *p*_trial 1_ = 0.048, *p*_trial 2_ = 0.001, *p*_trial 3_ = 0.03, *p*_trial 4_ = 0.03) (Fig. 1B, Conditioning).

**Figure 1.**
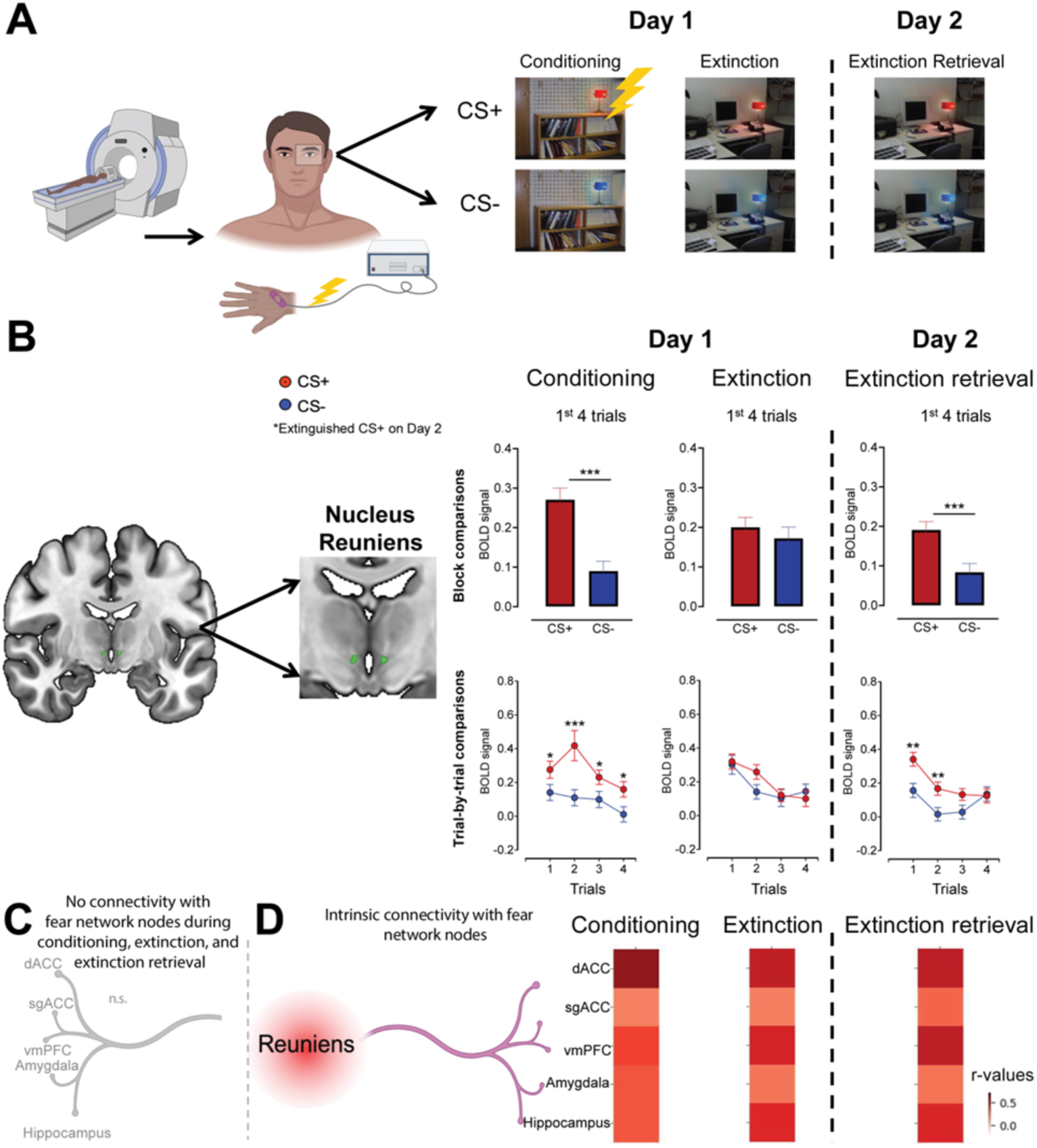
Human RE encodes aversive CSs during fear conditioning and extinction. **(A)** Schematic of experimental protocol, in which human participants underwent conditioning and extinction on Day 1 and extinction retrieval on Day 2, while in an MRI scanner. **(B)** Coronal section showing the RE in the human brain (left) and BOLD signal in response to CS+ (red circles) and CS-(blue circles) across conditioning, extinction, and extinction retrieval (right). BOLD signal to CS+ and CS-is compared by averaging all 4 trials (top, block comparisons) as well as by a trial-by-trial comparison (bottom). **(C)** No functional connectivity was observed between RE and fear network nodes during conditioning, extinction, and extinction retrieval, despite positive significant intrinsic connectivity **(D)**. Data are represented as mean ± SEM. Some graphical elements in panels A, C, and D were created using BioRen-der (BioRender.com).

During extinction learning, we found no difference between the overall signal to the CS+ and CS-[*t*_319_ = 0.84, *p* = 0.404, mean difference 95 % CI (-0.04 - 0.09), Cohen’s *d* = 0.05 with 95 % CI (-0.06 - 0.16)]. The trial level analysis also showed no significant differences except a decrease in the signal across trials to both CS+ and CS-[main effect of Trial: *F*_2.83,876.63_ = 8.79, *p* < 0.001, partial *η*² = 0.028; no main effect of CS: *F*_1,310_ = 0.66, *p* = 0.42, partial *η*² = 0.002; no Trial x CS interaction: *F*_3,930_ = 1.12, *p* = 0.34, partial *η*² = 0.004. Post-hoc_FDR corrected_ for CS+ > CS-: *p*_trial 1_ to *p*_trial 4_ = 0.22 - 0.77] (Fig. 1B, Extinction).

In the extinction retrieval phase, our results pointed to a general heightened BOLD signal in response to CS+ compared to CS-[*t*_411_ = 4.07, *p* < 0.001, mean difference 95% CI (0.05 - 0.16), Cohen’s *d* = 0.20, 95 % CI (0.1 - 0.3)]. However, this overall difference was mainly derived from the very early trials of the retrieval test (first 2 trials) (main effect of Trial: *F*_2.9,1208.17_ = 7.27, *p* < 0.001; main effect of CS: *F*_1,411_ = 16.55, *p* < 0.001; Trial x CS interaction: *F*_2.93,1204.02_ = 2.48, *p* = 0.06. Post-hoc_FDR corrected_ for CS+ > CS-: *p*_trial 1_ = 0.004, *p*_trial 2_ = 0.008, *p*_trial 3_ = 0.053, *p*_trial 4_ = 0.84) (Fig. 1B, Extinction Retrieval).

To explore the role of the RE in coordinating cortical-hippocampal network activity, we examined the connectivity of the RE with brain regions associated with processing fear. The connectivity analysis revealed positive significant intrinsic connectivity of the RE with the dorsal anterior cingulate cortex (dACC), subgenual ACC (sgACC), ventromedial prefrontal cortex (vmPFC), amygdala, and HPC (Fig. 1D). However, we found that this connectivity is decreased when the RE is engaged in an actual fear learning process, insofar as we found no significant connectivity values with any of the main ‘fear network’ regions during conditioning, extinction, and extinction retrieval (Fig. 1C).

These results reveal that the RE in humans encodes the associative value of an aversive CS, revealing heightened BOLD signal with conditioning and lowered signal during extinction and extinction retrieval. In addition, the connectivity findings reflect different neural mechanisms of the RE during resting states vs. fear learning. The RE might suppress the default connectivity to play a more localized role that focuses on processing fear rather than communicating the information to other brain regions.

These results suggest that the RE in humans encodes perceived salience and associative value of the conditioned stimulus, revealing heightened BOLD signal with conditioning and lowered signal during extinction and extinction retrieval. In addition, the connectivity findings reflect different neural mechanisms of the RE during resting states vs. fear learning. The RE might suppress the default connectivity to play a more localized role that focuses on processing fear rather than communicating the information to other brain regions.

### Calcium activity in the rat RE encodes associative value during fear conditioning and extinction

To determine if RE activity in rodents shows similar learning-related changes as in humans, we recorded calcium transients from the RE using fiber photometry (Fig. 2A). Recordings took place as adult male (*n* = 4) and female (*n* = 4) rats underwent habituation, auditory fear conditioning, context extinction and retrieval, and cued extinction and retrieval (Fig. 2C). Male and female data were collapsed because no significant sex differences were found.

**Figure 2.**
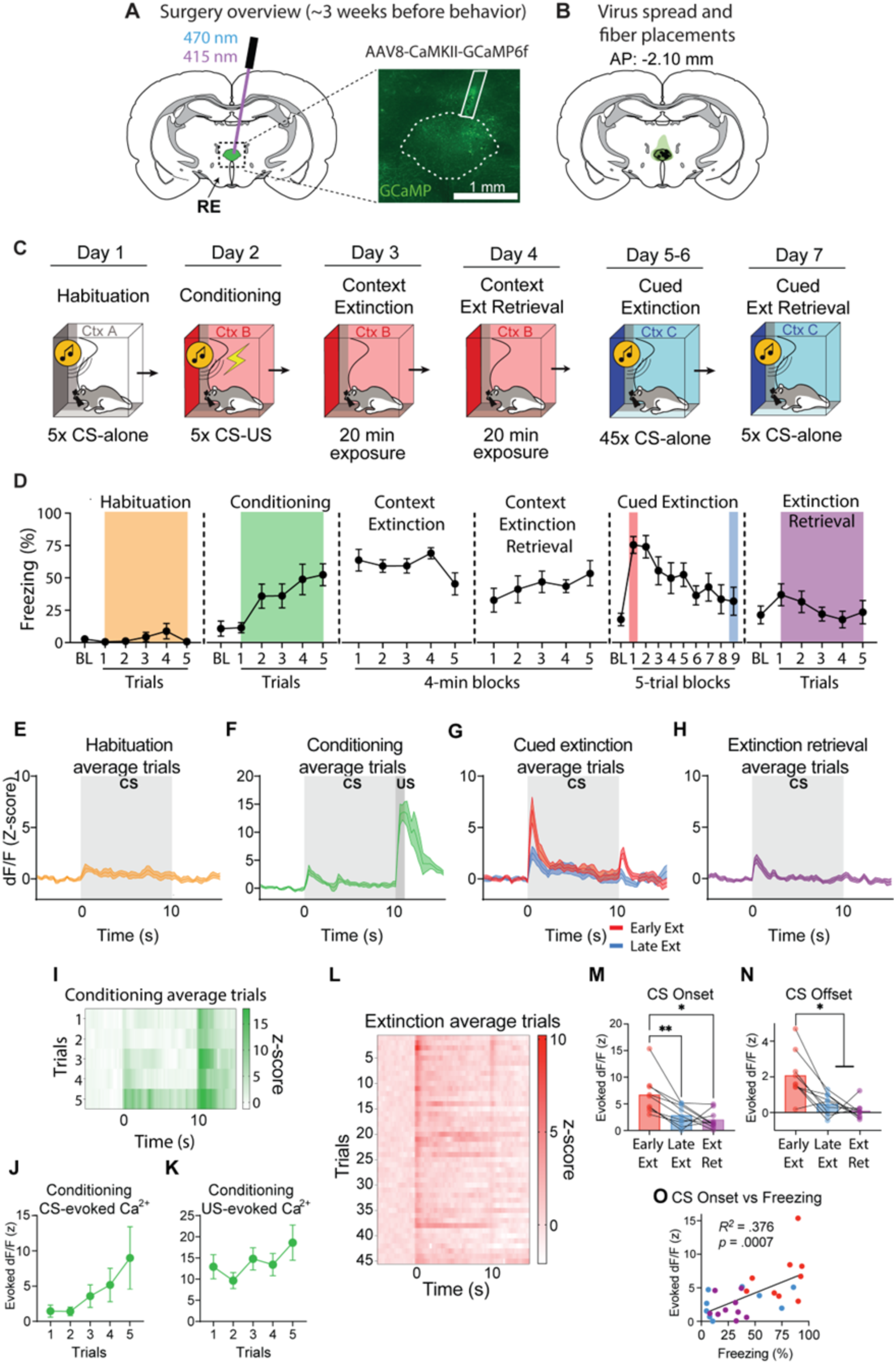
Calcium activity in the rat encodes associative value during fear conditioning and extinction. **(A)** Adeno-associated viruses (AAVs) encoding the GCaMP6f fluorescent calcium indicator were injected into the RE approximately three weeks before behavioral experimentation and recordings. GCaMP fluorescence was recorded with a chronically implanted fiber optic by interleaving blue (470 nm; GCaMP) and violet (410 nm; isosbestic) light. **(B)** Representative viral spread and fiber tip locations for all rats (*n* = 8). **(C)** Experimental timeline. **(D)** Percentage of freezing behavior levels during each of the recording days averaged for all animals. CS-evoked fluorescence averaged across trials and animals during habituation **(E)**, conditioning **(F)**, cued extinction **(G)**, and extinction retrieval **(H)**. **(I)** Heatmap showing averaged CS-evoked activity across animals for all five trials of conditioning. **(J)** CS-and **(K)** US-evoked activity across trials during conditioning. **(L)** Heatmap showing averaged CS-evoked activity across animals for all 45 trials of extinction. Average peak evoked fluorescence at CS Onset **(M)** and CS Offset **(N)** during early extinction, late extinction, and extinction retrieval. **(O)** Linear regression showing that average peak fluorescence at CS onset across early extinction, late extinction, and extinction retrieval positively correlates with freezing behavior. Data are represented as mean ± SEM.

During habituation, the RE was moderately responsive to the novel CS (Fig. 2E, *t*_8_ = 4.688, *p* =.0016), despite all animals exhibiting low freezing (no main effect of Trials: *F*_5,35_ = 1.171, *p* =.34) (Fig. 2D, Habituation). RE activity increased across conditioning trials as animals acquired fear of the CS (main effect of Trials: *F*_5,35_ = 5.062, *p* =.0014) (Fig. 2D, Conditioning), though the increases in CS-evoked activity were not statistically significant (no main effect of Trials: *F*_1.418,7.092_ = 1.767, *p* =.2338) (Fig. 2F, I, J). The RE also showed strong US-evoked activity during the first conditioning trial, which did not change significantly across trials (no main effect of Trials: *F*_4,20_ = 1.156, *p* =.3597) (Fig. 2K). A repeated-measures ANOVA on the CS-and US-evoked responses normalized to the percent of maximum response for each animal confirmed these observations (Stimulus x Trials interaction: *F*_4,59_ = 2.82, *p* =.033).

During the cued extinction session, all animals successfully extinguished fear to CS and exhibited a reduction in freezing across trials (main effect of Trial blocks: *F*_2,18_ = 6.408, *p* =.006) (Fig. 2D, Cued Extinction). CS-evoked calcium responses were maximal at the outset of extinction training and decreased over the course of extinction (Fig. 2G, L). To confirm this, we compared CS-evoked activity at the CS onset during early (first five) vs late (last five) extinction trials and observed a significant difference (main effect of Time: *F_2,14_* = 12.13, *p* =.0013; Early vs Late post-hoc: *p* =.0056) (Fig. 2M). The RE also displayed increased activity at the time of CS offset; this effect was evident during early extinction trials which diminished across extinction training (main effect of Time: *F_2,14_* = 12.72, *p* =.0013; Early vs Late post-hoc: *p* =.0102) (Fig. 2N) and may reflect a prediction error associated with omission of the US.

During extinction retrieval, all animals displayed low freezing levels, demonstrating successful extinction memory retrieval (no main effect of Trials: *F*_10,40_ = 1.17, *p* =.3365) (Fig. 2D Extinction Retrieval). We found that CS-evoked activity in the RE was also low (Fig. 2H) compared to early extinction for both CS onset (Tukey’s post-hoc: *p* =.0114) and CS offset (Tukey’s post-hoc: *p* =.0103); CS-evoked activity during the retrieval trials was like late extinction activity for both CS onset and offset (*p* >.05) (Fig. 2M,N). We additionally found that CS-evoked calcium activity in the RE strongly correlates with CS-evoked freezing behavior across extinction and extinction retrieval (Fig. 2O, *r*^2^ =.376, *p* =.0007). Taken together with our findings in humans, these data reveal that CS-evoked activity in the RE is strongly associated with conditioned fear.

### RE calcium activity tracks defensive freezing behavior

It was previously discovered that the RE plays a critical role in suppressing conditioned freezing behavior^39^, but it is unclear if neural activity in RE correlates with or predicts defensive freezing. To determine this, we analyzed spontaneous activity in the RE during exposure to the conditioning context prior to conditioning (context habituation), after conditioning (context extinction), and after context extinction (context retrieval). Animals showed high freezing levels during context extinction and context retrieval sessions compared to habituation (mixed model main effect: *p* <.0001; Hab vs Ext post-hoc: *p* <.0001, Hab vs Ret post-hoc: *p* =.0018). Average freezing levels across these sessions decreased from context extinction to context retrieval session, though this decrease was not significant (Ext vs Ret post-hoc: *p* <.0644) (Figs. 2D&3A).

We first looked at potential differences in RE activity between bouts of locomotor activity and freezing during the habituation session to determine if RE activity tracks general activity (Fig 3B-D). We found no significant changes in RE activity between bouts of activity and immobility during the habituation session (Fig. 3C-D, Start vs Stop: *t_5_* = 0.819, *p* =.45), demonstrating that the RE does not track nonspecific changes in locomotor activity. However, during both context extinction (Fig. 3F-H) and retrieval (Fig. 3J-L), we found that calcium activity in the RE correlates with defensive freezing. Specifically, initiation of freezing behavior was associated with a marked decrease in RE activity (Fig. 3H&L, Extinction: Start: *t_8_* = 4.239, *p* =.0028; Retrieval: Start: *t*_6_ = 4.099, *p* =.0064), whereas termination of freezing was associated with an increase in RE activity (Fig. 3H&L, Extinction: Stop: *t_8_* = 6.254, *p* =.0002; Retrieval: Stop: *t*_6_ = 6.858, *p* =.0005). In line with previous work, we used the third derivative of smoothed and averaged calcium traces^40^ (Fig. 3M) and found that the increase in RE activity began ∼500 ms prior to freezing termination (Fig. 3N, Extinction: *t_7_* = 11.67, *p* <.0001; Retrieval: *t_6_* = 7.767, *p* =.0002), suggesting a role for RE activity in actively suppressing freezing behavior. Although there was no difference between extinction and retrieval days on freezing driven RE activation (Fig. 3E, Ext vs Ret post-hoc: *p* =.8173) or suppression (Fig. 3I, Ext vs Ret post-hoc: *p* =.7503), increases in RE activity occurred in closer temporal proximity to freezing termination during retrieval compared to extinction (Fig. 3N; *t*_5_ = 2.795, *p* =.0382). Collectively, these findings in rodents suggest that the RE displays two distinct profiles of activity, with one encoding the associative value of shock-predictive cues, and the other encoding changes in defensive freezing that occur in shock-paired contexts. Interestingly, short-latency CS-evoked activity in RE correlates with high freezing, whereas context-associated activity correlates with low freezing.

**Figure 3.**
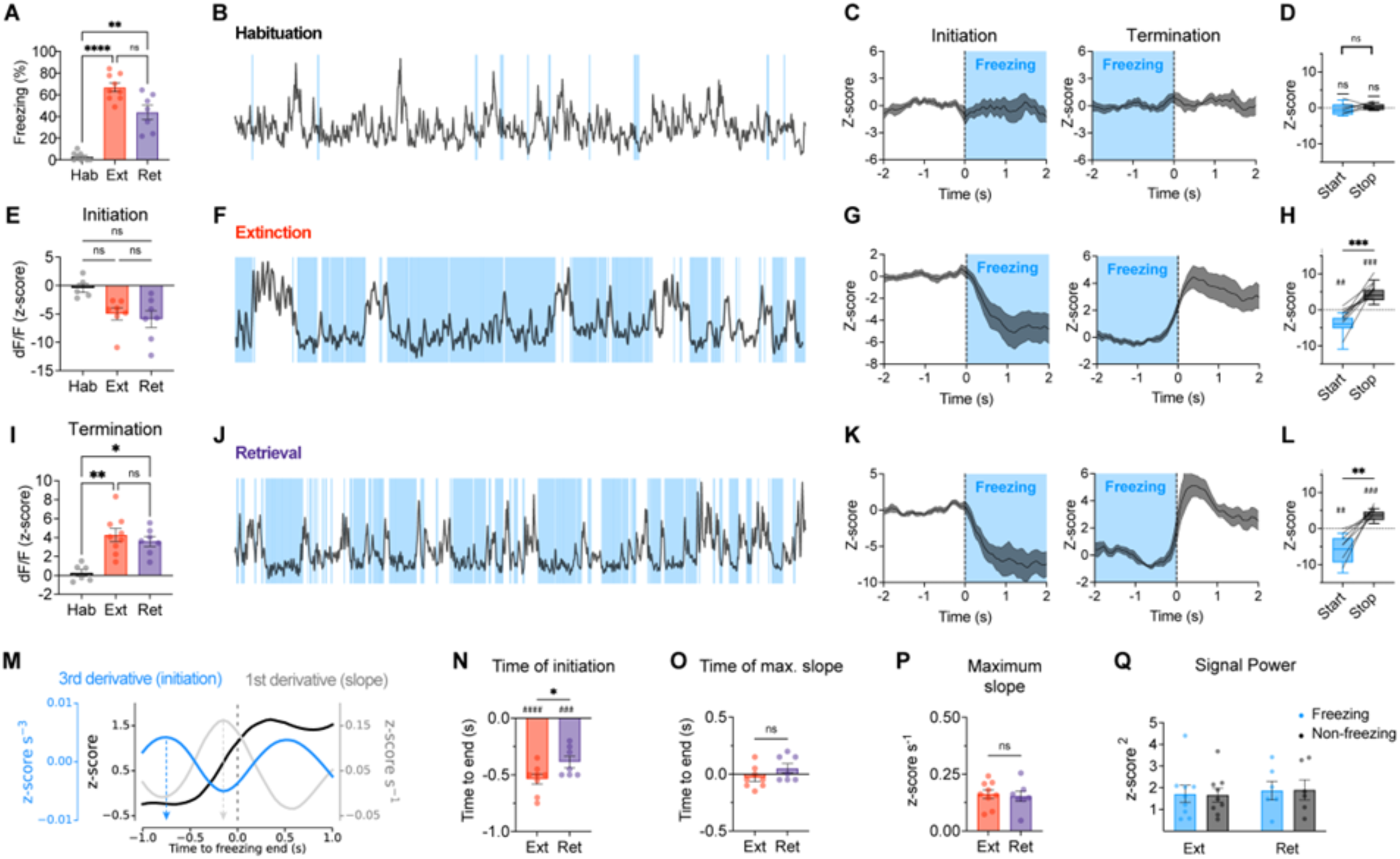
Spontaneous calcium activity in the RE encodes defensive state transitions in shock-paired contexts. **(A)** Average freezing behavior during habituation, context extinction, and context extinction retrieval. **(B,F,J)** Representative traces of RE activity during habituation, context extinction, and context retrieval, respectively, with freezing epochs denoted by blue shades. **(C,D)** Average RE activity during the initiation and termination of non-fear related immobility during habituation, compared to averaged RE activity during the initiation and termination of fear related freezing behavior during context extinction **(G,H)** and retrieval **(K,L)**. Average change in the RE activity during freezing initiation **(E)** and termination **(I)** across days. **(M)** Example graphic of the third derivative of a smoothed photometry trace for determining the timepoint of RE activation relative to freezing termination. Average time point of RE activation **(N)**, maximum slope **(O)**, and amplitude of maximum slope of the first derivative **(P)** prior to freezing termination for extinction and retrieval. **(Q)** Average signal power during freezing and non-freezing epochs during context extinction and retrieval. Data are represented as mean ± SEM.

### Single-unit activity in the RE during fear conditioning and extinction

In the previous experiments, averaging the BOLD and calcium signals within the RE provided a population measure of RE activity integrated across hundreds of neurons. To characterize RE activity at the level of single neurons, we performed extracellular single-unit recordings using the same experimental timeline as in fiber photometric recordings (Fig. 2C). We implanted animals with multi-electrode arrays (16-channel; *n* = 21, 11 male and 10 female or 32-channel; *n* = 3, 2 male and 1 female) in the RE. One animal was excluded from analyses due to missed target, resulting in *n* = 23 (13 male and 10 female).

Animals showed low freezing levels during habituation (Fig. 4C, Habituation) (no main effect of Trials: *F*_5,105_ = 0.39, *p* =.85), with no sex difference (*F*_1,21_ = 2.06, *p* =.17). Single-unit recordings yielded 85 recorded neurons during habituation, most of which remained unresponsive to the CSs (77/85) (Fig. 4D, Habituation). Animals acquired freezing to the CSs during conditioning (Fig. 4C Conditioning) (main effect of Trials: *F*_5,105_ = 37.86, *p* <.0001) with males showing overall higher freezing compared to females (main effect of Sex: *F*_1,21_ = 14.20, *p* =.0011) yet similar rates of learning (no Trials x Sex interaction: *F*_5,105_ = 1.74, *p* =.131). A total of 110 neurons were recorded during conditioning. Of these neurons, 72 responded to either the CS, US, or both. Eight were exclusively responsive to the CS (≥3 *z*-score, see Methods), 47 were responsive only to the US, 17 were responsive to both the CS and US, and 38 were unresponsive. Of the responsive neurons, this corresponds to 11%, 65%, and 24% for CS-excited, US-excited, and CS-and US-excited neurons, respectively (Fig. 4E). Average normalized firing of the CS-responsive neurons (25/110) revealed increased firing with the CS and US presentations, with no significant difference between the two (Wilcoxon signed rank test, *p* =.1771) (Fig. 4D, Conditioning).

**Figure 4.**
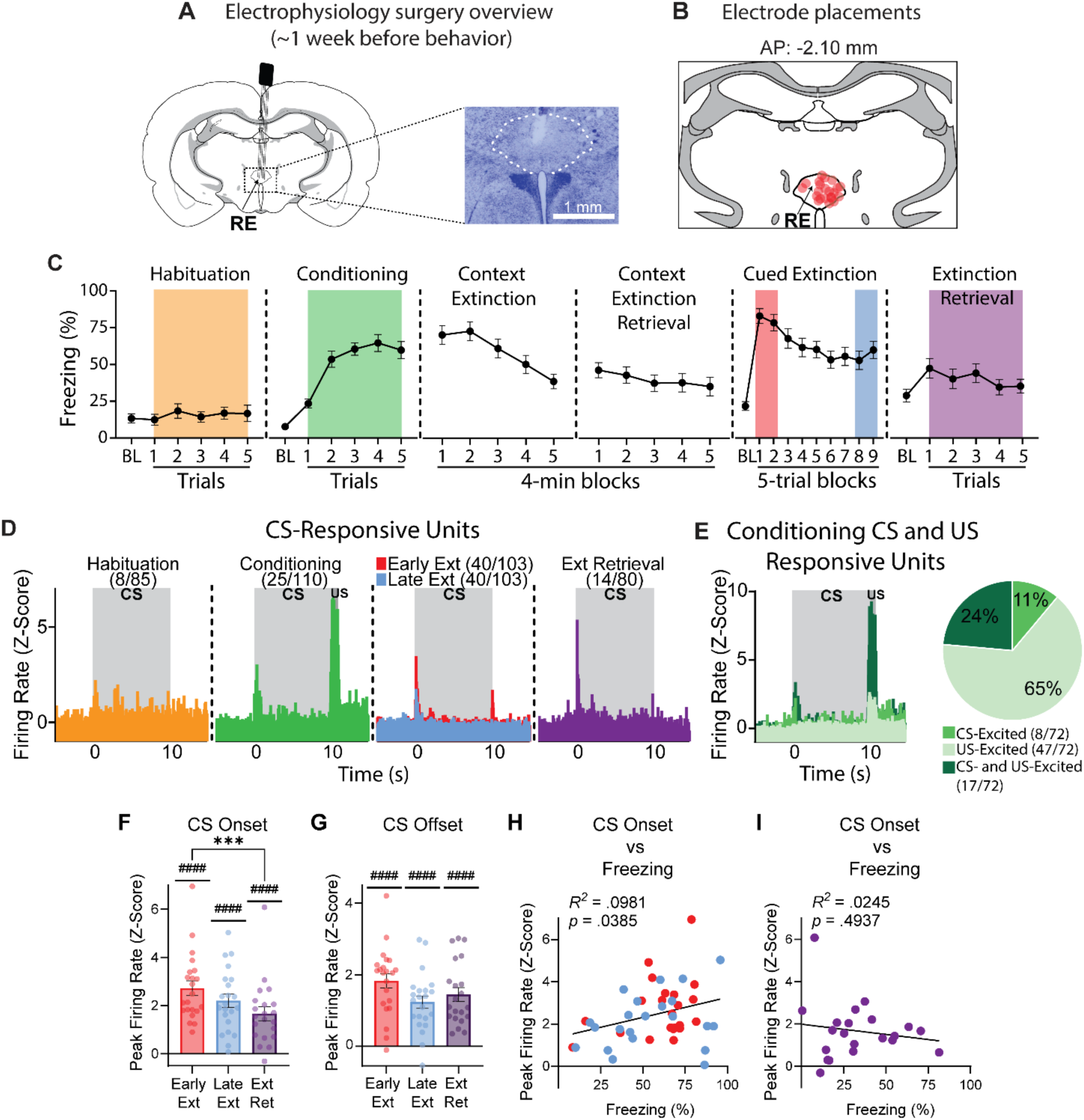
Neurons in the RE encode associative value during fear conditioning and extinction. **(A)** Microelectrode arrays were chronically implanted in the RE approximately one week before behavioral experimentation and recording. **(B)** Electrode placements based on electrode tips in all animals (*n* = 23). **(C)** % freezing behavior levels during each of the recording days averaged for all animals. **(D)** Average normalized firing rates of CS-responsive RE neurons during habituation, conditioning, extinction, and extinction retrieval. **(E)** Average normalized firing rates of CS-excited, US-excited, and CS-and US-excited RE neurons during conditioning (left) and the pie chart showing their proportions (right). Average peak evoked firing at CS Onset **(F)** and CS Offset **(G)** during early extinction, late extinction, and extinction retrieval. **(H)** Linear regression showing that peak evoked firing at CS onset during early extinction and late extinction positively correlates with freezing behavior. **(I)** Linear regression of peak evoked firing at CS onset during extinction retrieval vs freezing reveals a negative slope. Data are represented as mean ± SEM.

During cued extinction (Fig. 4C), animals extinguished fear to the CS as evidenced by robust decrease in freezing across trials (main effect of Trial blocks: *F*_9,189_ = 23.60, *p* <.0001) (Fig. 4C Cued Extinction). There was no sex difference (*F*_1,21_ = 3.86, *p* =.063) or Trial blocks x Sex interaction (*F*_9,189_ = 1.135, *p* =.3397). Extinction recordings yielded 103 neurons, 40 of which were CS-responsive. Confirming fiber photometric recordings, CS responsiveness was greater at CS-onset during early extinction (first 10) compared to late extinction (last 10) (Wilcoxon signed rank test, *p* =.0006). A more detailed analysis of the CS-onset response latencies revealed that CS-responsive neurons, on average, exhibited increases in activity roughly 70 ms after CS onset, with significantly higher firing rates between 70-90 ms (Wilcoxon signed rank test, *p*s <.05). This suggests that a multisynaptic pathway conveys auditory information to the RE. Earlier findings from the lateral amygdala neurons demonstrated much shorter CS-response latencies (< 20 ms), in line with monosynaptic projections from the auditory thalamus to the amygdala^41,42^. Longer response latencies of RE neurons, however, may involve a thalamo-amygdala-RE pathway, consistent with direct amygdala-to-RE projections^22^.

During extinction training, CS responsive neurons also showed increased firing at CS offset during early extinction, which diminished across extinction training (Wilcoxon signed rank test, *p* <.0001, Fig. 4D, Early Ext vs Late Ext). Additionally, CS-evoked firing in the RE during early and late extinction was positively correlated with CS-evoked freezing (Fig. 4H, *r*^2^ =.0981, *p* =.0385), a finding that confirms the fiber photometry results. During extinction retrieval, animals exhibited substantially lower freezing compared to extinction training (*t*_22_ = 4.454, *p* =.0002). There was no sex difference (*F*_1,21_ = 0.003, *p* =.9567) or Trials x Sex interaction (*F*_5,105_ = 0.3895, *p* =.855). Of the 80 neurons recorded during extinction retrieval, a small population was CS-responsive (14/80). These neurons fired strongly at the CS onset (Fig. 4D, Ext Retrieval). Finally, CS-evoked firing during extinction revealed a negative slope with freezing behavior, although not significant (Fig. 4I, *p* >.05). This is in contrast with the low calcium signal observed during extinction retrieval and suggests that firing of a smaller proportion of RE neurons may promote the suppression of fear during extinction retrieval.

Because CS-responsive neurons consist of a smaller proportion of all recorded neurons, we also examined CS-evoked firing in all neurons, averaged within animals. Comparing the CS onset firing during early extinction, late extinction, and extinction retrieval, we observed decreased firing over time (main effect of Time: *F*_2,40_ = 8.76, *p* =.0007). CS onset firing was significantly higher during early extinction compared to extinction retrieval (Early Ext vs Ext Ret post-hoc: *p* =.004) (Fig. 4F). CS offset firing difference between early extinction, late extinction, and extinction retrieval just missed significance (main effect of Time: *F*_2,40_ = 2.875, *p* =.0681) (Fig. 4G).

Overall, as in the population measures of RE activity in humans and rats, conditioning was characterized by strong CS-evoked single-unit activity which weakened with extinction learning, suggesting that the RE neurons encode the associative value of the CS. Despite overall decreased firing, single-unit recordings revealed a subpopulation of RE neurons that fire strongly to CSs during extinction retrieval. This suggests that RE CS-evoked firing predicts fear suppression in addition to fear expression.

### RE neurons display heterogeneous firing patterns during extinction

On average, RE neurons show robust increases in CS-evoked activity after conditioning that decrease over the course of extinction. Here we aimed to determine if individual RE neurons within the population show divergent patterns of activity. First, we sought to answer if fear and extinction memories are encoded by distinct ensembles in the RE. We classified neurons into subpopulations based on their early vs late extinction firing using the following criteria: ‘nonresponsive’ if there was no CS-evoked response (*z*-score < 3) during early and late extinction trials, ‘fear non-decrement’ if there was a significant CS-evoked response (*z*-score ≥ 3) during both early and late extinction trials, ‘fear’ if there was a CS-evoked response during early extinction trials but not late extinction trials, and ‘extinction’ if there was a CS-evoked response during late extinction trials but not early extinction trials. This resulted in the following categories: nonresponsive cells (46/103), fear non-decrement cells (19/103), fear cells (21/103), and extinction cells (17/103) (Fig. 5A). Of the CS-responsive cells, this corresponds to 33%, 37%, and 30% for fear non-decrement cells, fear cells, and extinction cells, respectively (Fig. 5B). These different cell types revealed a relatively homogeneous distribution among animals (Fig. 5C), with some animals showing all the cell types. We additionally performed hierarchical clustering on normalized firing from each unit, aligned to the CS (CS onset or offset) during early extinction and late extinction trials, separately. These analyses revealed three distinct CS-evoked firing patterns during both early and late extinction: increase, decrease, and offset change (see Fig. S1 for details). These results demonstrate that RE neurons respond to CSs similarly during both early and late extinction. The majority of the RE neurons show matched CS onset and offset responses (both decrease or both increase in firing), with a third population of neurons showing no change in firing during CS onset but offset.

**Figure 5.**
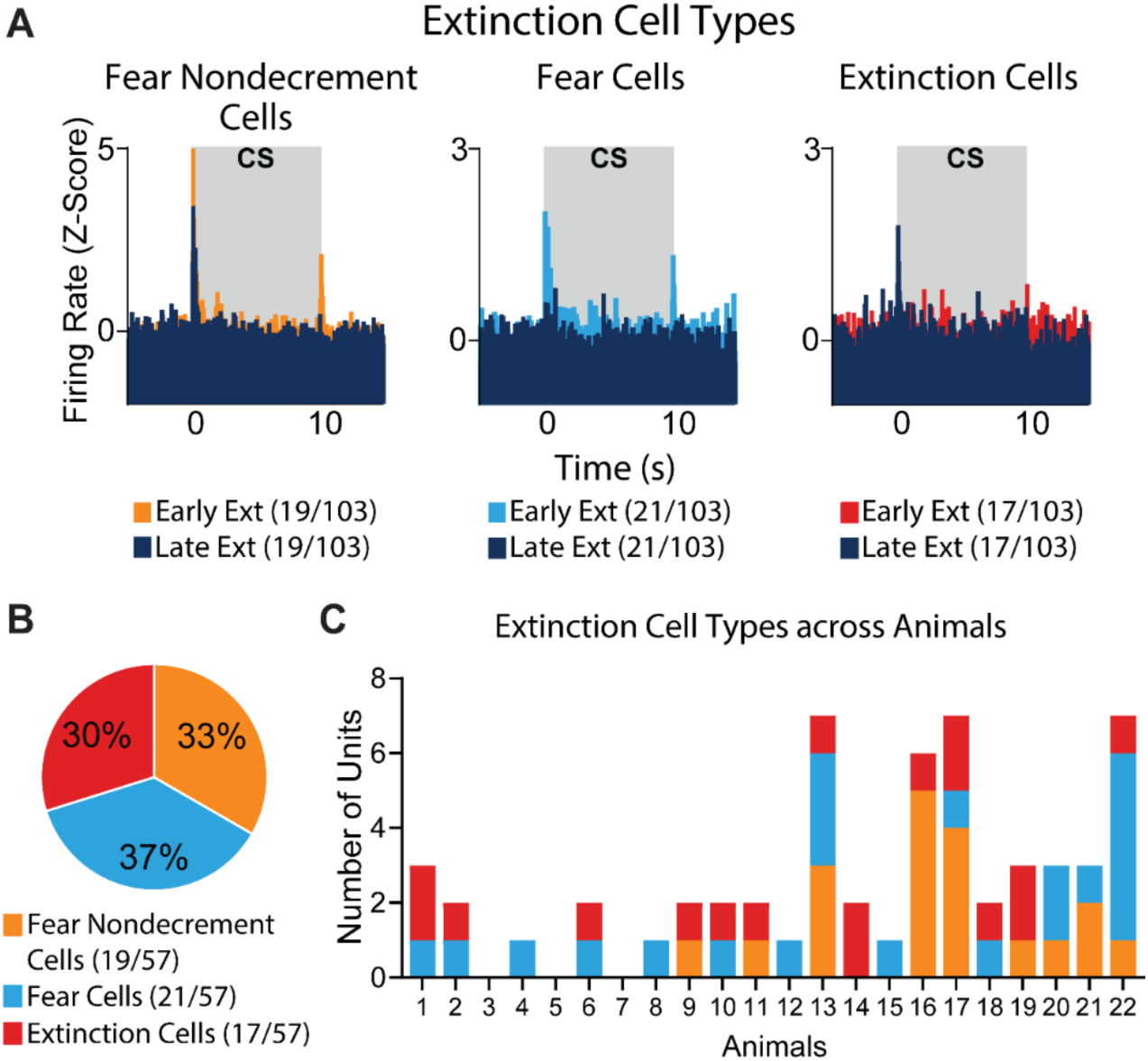
RE neurons display heterogenous firing patterns during extinction. **(A)** Average normalized firing rates of RE neuronal subpopulations with different CS-firing patterns during extinction: fear non-decrement cells (left), fear cells (middle), and extinction cells (right). **(B)** Pie chart showing the proportions of the cell types in **(A)**. **(C)** Stacked bar plot showing the distribution of different cell types across animals.

Overall, RE single-unit recordings reveal that there is considerable heterogeneity in RE neuronal activity to CSs during extinction. Distinct ensembles encode fear versus extinction memories. Fear cells and extinction cells show the opposite firing pattern with the former firing during early extinction and the latter during late extinction. “Extinction” neurons in the RE may play an important role in suppressing conditioned fear responses.

### Spontaneous firing of RE neurons tracks defensive freezing states

Fiber photometric recordings revealed that the spontaneous fluctuations in RE calcium activity are highly correlated with conditioned freezing behavior and increases in RE activity reliably precedes transitions from freezing to activity. Here, we examined if RE single-unit activity similarly covaries with freezing and movement. For this, we similarly focused on context extinction and retrieval recording days.

Animals showed high freezing levels during context extinction which decreased over time (main effect of Trials: *F*_4,84_ = 16.14, *p* <.0001) and remained low during context retrieval (no main effect of Trials: *F*_4,84_ = 1.47, *p* =.2198) (Fig. 4C, Context Extinction and Context Extinction Retrieval). The main effect of Sex or Trials x Sex interaction for both days was not significant (*p*s >.05). For the subsequent analyses, we analyzed data from both context extinction and retrieval days. Because we found similar results, we present data only from context retrieval [*n* = 131 units; see Fig. S2 for context extinction (*n* = 115 units)]. We averaged normalized firing from each unit for a-1 s to 1 s period, aligned at behavioral transition (freezing initiation or termination). Freezing initiation was associated with a decrease in firing (Fig. 6A, Initiation, Wilcoxon signed rank test, *p* <.0001), with the decrease in firing preceding the freezing initiation. We also observed an increase in firing with freezing termination, although not significant (Fig. 6A, Termination, *t*_39_ = 0.3323, *p* =.7414). We then averaged units within animals creating a single value for each animal. A two-way repeated measures ANOVA with Non-freezing v Freezing and Initiation v Termination (matched for both factors) confirmed that firing is significantly lower during Freezing compared to Non-freezing bouts (main effect of Non-freezing v Freezing, Fig. 6B, *F*_1,21_ = 143.9, *p* <.0001; Post-hocs for Initiation: Freezing vs Non-freezing, Termination: Freezing vs Non-freezing, Freezing: Initiation vs Non-freezing: Termination, Freezing: Termination vs Nonfreezing: Initiation: *p*s <.0001). This confirmed the previous fiber photometry results, suggesting that calcium transients from the RE reflect neural spiking.

**Figure 6.**
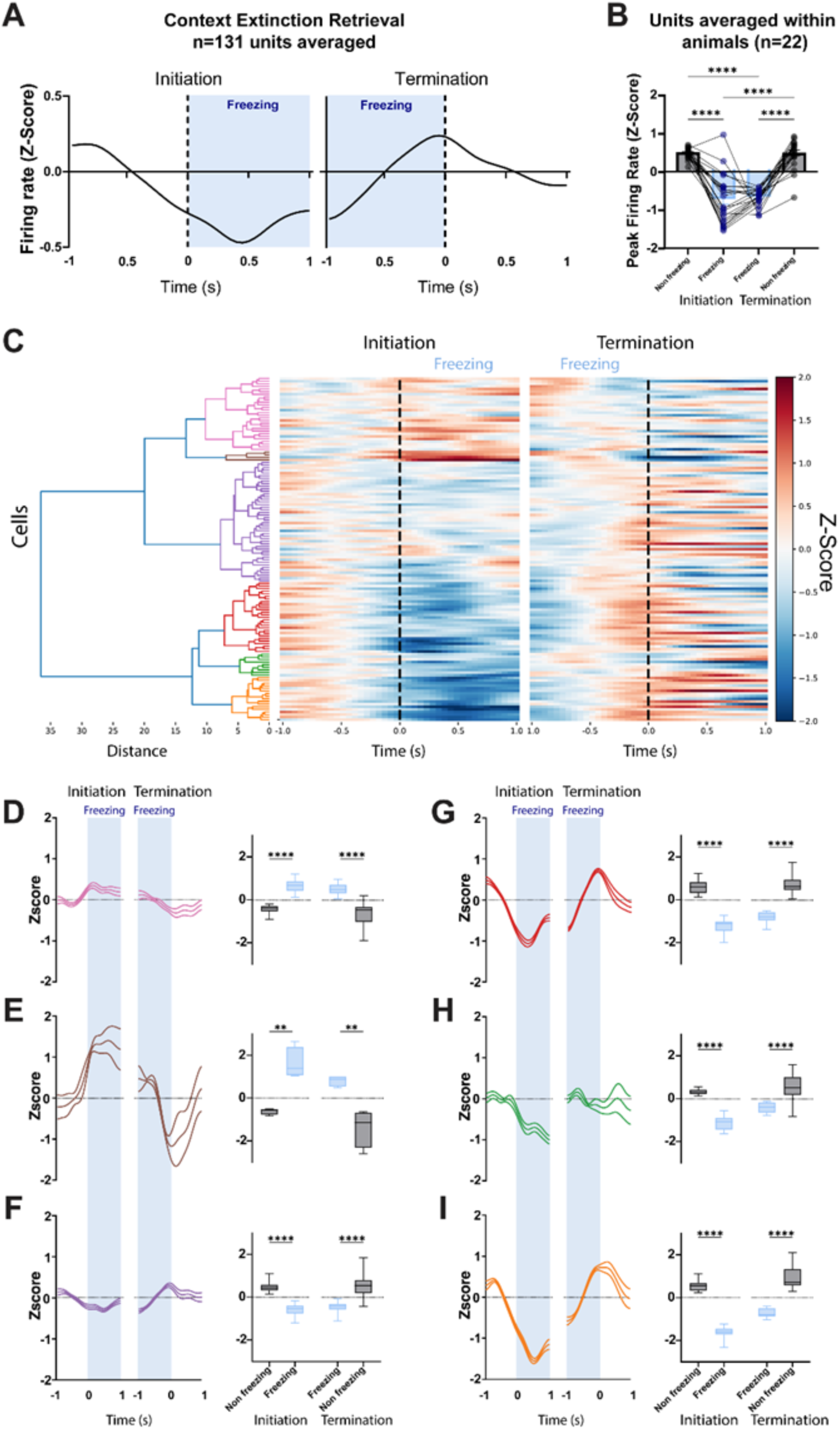
Spontaneous firing of RE neurons tracks behavioral state transitions. **(A)** Average normalized firing of neurons (*n* = 131) over time during freezing initiation (left) and termination (right). **(B)** Bar graph with peak normalized firing of neurons in nonfreezing and freezing epochs during freezing initiation and termination. **(C)** Dendrogram showing six different clusters based on hierarchical clustering of normalized firing over time. **(D-I)** Average normalized firing of each cluster over time (left) and box and whisker plots (right) showing peak normalized firing in non-freezing and freezing epochs during freezing initiation and termination. Data are represented as mean ± SEM.

Despite the dominant pattern in which RE firing decreases with freezing, we sought to examine neuronal responses in more detail. To this end, we performed hierarchical clustering on normalized firing from each unit, aligned at behavioral transition (freezing initiation or termination). This yielded six distinct clusters (Fig. 6C-I). Among these clusters, Cluster 1 and 2 neurons revealed increased firing with freezing as opposed to the dominant pattern (Fig. 6D-E). A two-way repeated measures ANOVA confirmed significantly higher firing for Freezing compared to Non-freezing bouts during both freezing initiation and termination (main effects of Non-freezing v Freezing, Fig. 6D right, Cluster 1: *F*_1,27_ = 230, *p* <.0001, Post-hocs for Initiation: Freezing vs Non-freezing, Termination: Freezing vs Non-freezing: *p*s <.0001; Fig. 6E right, Cluster 2: *F*_1,3_ = 49.96, *p* =.0058, Post-hocs for Initiation: Freezing vs Non-freezing, Termination: Freezing vs Non-freezing: *p*s <.002). On the other hand, Cluster 3-6 neurons revealed decreased firing with freezing (Fig. 6F-I), the pattern observed with average normalized firing of all units (Fig. 6A) and fiber photometric recordings (Fig. 3). A similar two-way repeated measures ANOVA confirmed that firing is significantly lower during Freezing compared to Non-freezing for all clusters (main effects of Non-freezing v Freezing, Fig. 6F right, Cluster 3: *F*_1,45_ = 730.4, *p* <.0001; Fig. 6G right, Cluster 4: *F*_1,26_ = 914.6, *p* <.0001; Fig. 6H right, Cluster 5: *F*_1,8_ = 128.7, *p* <.0001; Fig. 6I right, Cluster 6: *F*_1,16_ = 649.5, *p* <.0001, Post-hocs for Initiation: Freezing vs Non-freezing, Termination: Freezing vs Non-freezing: *p*s <.0001).

Overall, these data show that spontaneous single-unit firing in the RE, like spontaneous fluctuations in its calcium activity, is highly correlated with freezing behavior after context extinction; spike firing in the majority of RE neurons reliably increases prior to transitions from freezing to activity. In addition, a smaller proportion of RE neurons show the opposite pattern (decrease in firing from freezing to activity). These results confirm the involvement of the RE in the regulation of defensive states.

## DISCUSSION

Here we demonstrate that the RE has a conserved role in fear and extinction memory processes across rats and humans. Fear conditioning in both rats and humans is associated with an increase in CS-evoked activity in the RE. Conditioned increases in RE activity decreases after extinction learning and remains low during extinction retrieval. Single-unit recordings from rodents also revealed that the RE modulates defensive state transitions. Importantly, these data demonstrate that the RE has two separate representations of associative fear. First, cue-evoked RE activity encodes the associative value of an aversive CS, evidenced by increases in short-latency CS-evoked BOLD signal, calcium transients, and single-unit discharges. In these cases, high levels of RE activity are associated with the expression of conditioned fear. Second, the RE encodes a state signal, evidenced by fluctuations in spontaneous activity that track defensive behavior and are inversely related to conditioned freezing.

Both the human and rodent thalamus have been proposed to do more than simply relay information between cortical and subcortical structures, including contextual modulation of cortical representations^43–46^. Within each thalamic nucleus, distinct microcircuits may exhibit distinct functions, depending on their inputs and outputs^44^. However, these functions and their conservation across species are largely unknown. The RE, with its bidirectional projections with the mPFC and HPC, is well-positioned to affect both cortical and hippocampal processes, including memory. Data from the human RE have been scarce, mostly due to the small size of the RE in the human brain. However, similar to the rodent brain, the RE is believed to function as a hub by coordinating PFC-HPC interactions in the human brain^12,31,47^. The present results are the first in showing the conserved role of the RE in humans and rodents in processing emotional memories.

One way the RE might mediate memory retrieval after extinction is through “retrieval suppression’’, a process hypothesized to prevent the retrieval of a context-inappropriate fear memory when the extinguished CS is encountered^24^. The RE may relay mPFC input to the HPC to suppress the retrieval of fear memories and to retrieve extinction memories^30^. Medial prefrontal cortical input to RE neurons during extinction retrieval may underlie increased firing to CSs. The RE, then, projects to the HPC to mediate retrieval suppression^24–26^. Consistent with this model, retrieval suppression in humans, is associated with activation in the right dorsolateral and ventrolateral PFC that is accompanied by reductions in HPC activity^24^. Data from rodents suggest a direct role for the RE in retrieval suppression, as inhibiting the RE or its projections with the mPFC and HPC impairs extinction learning and retrieval^30,34–36^.

However, some aspects of the present data may seem at odds with the retrieval suppression model, insofar as RE did not show functional connectivity with the vmPFC and HPC during conditioning, extinction, and retrieval in humans. However, it should be noted that a lack of connectivity may not necessarily mean independence^48,49^. Indeed, previously, we showed mPFC-HPC oscillatory synchrony during extinction retrieval in rats, which was impaired by RE inactivation^36^. Another aspect at odds with the retrieval suppression model is that recordings in both humans and rodents revealed lowered activity in the RE after extinction learning and during extinction retrieval. These data suggest that CS-evoked responses in the RE predict fear expression rather than fear suppression. However, single-unit recordings in rats revealed a subpopulation of RE neurons that show increased firing only in the later trials of extinction (extinction cells). Extinction memory may be encoded by this ensemble, which may also show increased firing during extinction retrieval. Indeed, despite overall decreased firing, CS-responsive cells showed strong firing to the CSs during extinction retrieval.

We previously showed that a small number of RE neurons displays increased firing to extinguished CSs during extinction retrieval, but not fear renewal^30^, suggesting that RE neuronal firing may be necessary for the suppression of fear and retrieval of extinction memories. Similarly, we observed an increased RE BOLD signal to CS+ during extinction retrieval, a difference derived from the first two trials. In the last two trials, this signal was comparable to the CS-signal. Single-unit recordings during retrieval also support previous findings^30^, as a small subpopulation of the recorded cells showed strong firing to CSs. This CS-evoked firing during extinction retrieval revealed a negative slope with freezing behavior, although not significant. These findings provide further support that RE activity and neuronal firing during retrieval may be necessary to suppress fear in both humans and rodents.

The present results revealed that the RE, like the amygdala^50–52^ and the mPFC^50,53^, contains a heterogeneous population of neurons that separately encode fear (fear cells) and extinction (extinction cells). Extinction cells, constituting similar percentages as fear cells, may be responsible for suppressing conditioned fear. Extinction retrieval may involve retrieving extinction engrams and/or suppressing fear engrams in the HPC^54^. In line with this, recent work points to the somatostatin-positive interneurons in the ventral HPC for the extinction of contextual fear conditioning^55^. The RE may be targeting this subpopulation for the recall of extinction memories. In addition, the RE is involved in encoding the extinction context, necessary for the recall of context-dependent extinction memories^28,33^.

There is evidence that the RE may be directly responsible for the cessation of freezing behavior^34,39,56^. Here, we showed that the rodent RE encodes a state signal that negatively correlates with conditioned freezing. In the extinction context, RE calcium activity is lower when conditioned freezing is high and vice versa. Moreover, we observed the same pattern in RE single units, suggesting that calcium transients from the RE reflect neural spiking. These results are supported by previous results by Silva and colleagues^39^. Ratigan and colleagues^34^, on the other hand, reported the opposite pattern of activity (increased activity with freezing) recording calcium transients from RE axons in the dorsal CA1. Performing hierarchical clustering of single neuron responses, we further revealed clusters of neurons that show this opposite pattern (increased firing with freezing). Based on this, we speculate that the RE neurons displaying increased firing during freezing bouts may preferentially project to the HPC.

Overall, the RE encodes the associative value of aversive CSs and this activity parallels fear expression during conditioning and early trials of extinction. In addition, spontaneous fluctuations in RE activity correlate with fear suppression during late trials of extinction, extinction retrieval and in the extinguished fear context. Together with its strong connections with the mPFC, HPC, and other limbic structures including the amygdala^21–23^, the RE contributes to both fear and extinction memory processes in rats and humans. Fear expression and suppression roles of the RE may involve distinct circuitries, with the former possibly involving the amygdala and the latter involving the mPFC and HPC. While uncovering these distinct circuits awaits future research, we believe that the RE may be an important therapeutic target for fear, anxiety, and trauma-and stressor-related disorders, including the PTSD.

## METHODS

### Human subjects

The subjects were 293-412 healthy controls between 18 and 70 years old (M = 32.17 ± 13.1, ∼184 females). All participants were proficient in English, right-handed, and had normal or corrected-to-normal vision. The exclusion criteria included a history of seizures or significant head trauma, current substance abuse or dependence, metal implants, pregnancy, breastfeeding, or positive urine toxicology screen for drugs of abuse. We followed the latest version of Helsinki’s declaration, and all procedures were approved by the Partners HealthCare Institute Review Board of the Massachusetts General Hospital, Harvard Medical School. All participants provided written informed consent before taking part in the study. Some results from this dataset have been published elsewhere with a different focus^38,57,58^. The current results are novel and have not been previously published.

### Experimental procedure

All participants underwent a validated two-day fear conditioning and extinction paradigm^10,37^ while in an MRI scanner (see Fig. 1A). On day 1, the participants underwent a fear conditioning learning phase in which a light (e.g., red) was paired with a 500 ms electric shock (CS+) with a partial reinforcement rate of 62.5 %. Another light (e.g., blue) was also presented but never paired with the shock (CS-). The duration of each trial was 6 s and a fixation screen was presented during the inter-trial intervals for 12-18 s (15 s on average). The shock level was determined during pre-experiment calibration; participants were guided to select a highly annoying but non-painful shock level. The conditioning phase was followed by extinction learning, where stimuli were repeatedly presented with no shock. On day 2, the participants underwent an extinction retrieval phase to assess the retention of the fear extinction memory. All stimuli were presented with no shock within the same context we used in extinction. Finally, in our paradigm, we pseudorandomized and counterbalanced the order of CS+ and CS-across phases and between subjects.

### MRI data acquisition and preprocessing

The neuroimaging data were acquired using two different MRI settings. The first setting was in a Trio 3T whole-body MRI scanner (Siemens Medical Systems, Iselin, New Jersey) using an 8-channel head coil. The functional data in this setting were acquired using a T2* weighted echo-planar pulse sequence with these parameters: TR = 3.0 s, TE = 30 ms, slice number = 45, voxel size = 3 × 3 × 3 mm^3^. The second setting was in the same Trio 3T scanner using a 32-channel head coil. A T2* weighted echo-planar pulse sequence was applied to obtain the functional images using: TR = 2.56 s, TE = 30 ms, slice number = 48, voxel size = 3 × 3 × 3 mm^3^. The anatomical images in both settings were collected using a T1-weighted MP-RAGE pulse sequence parcellated into 1 × 1 × 1 mm^3^ voxels. We used elastic bands affixed to the head coil device to reduce head motions. We preprocessed the data using fMRIPrep, version 20.0.2^59,60^, and applied the default pipeline to correct slice timing, realign the functional images, and coregistration. We also normalized the data into the Montreal Neurological Institute (MNI) space and smoothed with a 6-mm full-width half-maximum Gaussian kernel.

### Activation analyses

To estimate the BOLD signal to CS+ vs. CS-, we applied the least-squares-based generalized linear model (GLM) for each participant using Statistical Parametric Mapping (SPM) 12. We estimated the beta values for each voxel during all stimuli in the paradigm (overall 32 regressors for the CS+ and CS-, a regressor for the context, and the shock). We also included the six parameters of the head movements (x, y, z directions, and rotations) in the GLM. We then used the contrast maps from the first level analysis to estimate the variability of these maps across all subjects (group-level analysis). The contrast maps from the second level were used to extract the averaged values across the voxels within a predefined mask of the RE. These regional signals were used to compare the BOLD response during each trial of CS+ and CS-. We averaged the signal in the first 4 trials to compare blocks of CS+ vs. CSusing t-statistics and ran repeated measures ANOVA on trial-by-trial signals. False discovery rate (FDR) correction was applied.

### Connectivity analyses

To compute the connectivity values, we used the CONN functional connectivity toolbox (version 22. a) for the MathWorks MATLAB program^61^. We segmented the anatomical data into tissues of gray matter, white matter, and CSF. We then applied a standard denoising pipeline^62^ to the functional data using the CONN default confounding parameters and regressed potential confounding effects using a component-based noise correction method. We also applied bandpass frequency filtering of the BOLD time series between 0.008 Hz and 0.09 Hz.

*Task-related connectivity*. To evaluate the difference in the connectivity values during the CS+ vs. CS-, we conducted generalized psychophysiological interaction analyses (gPPI) on blocks of the CS+ vs. CS-(each block consisted of the first 8 trials). We defined the RE as a seed and five main brain regions associated with processing threat as targets: dorsal anterior cingulate cortex (dACC), subgenual ACC (sgACC), ventromedial prefrontal cortex (vmPFC), amygdala, and HPC. We then examined how the CS+ and CS-conditions modulate the BOLD signal within both the RE and each target region separately. The gPPI was defined with seed BOLD signals as physiological factors, boxcar signals characterizing each task condition convolved with an SPM canonical hemodynamic response function as psychological factors, and the product of the two as psychophysiological interaction terms. Functional connectivity changes across conditions were characterized by the Fisher-transformed correlation coefficient of the psychophysiological interaction terms in each model. At the group level, a GLM model was used to assess task-related connectivity changes across participants. Differences between condition-related connectivity were evaluated using paired t-tests, with applying false discovery rate (FDR) correction at p < 0.05.

*Intrinsic connectivity*. To estimate connectivity during the resting state, we performed seed-based connectivity analysis (SBC) on the resting-state time series, excluding task-related activity and defining a ‘rest’ condition^63^. For each subject, we computed the Fisher-transformed correlation coefficients between the RE and each targeted ROI using a weighted GLM. At the group level, we applied a second GLM to the Fisher Z-transformed connectivity maps across participants^62^. To assess whether the average connectivity differed from zero, t-statistics were computed for each ROI-to-ROI pair. False discovery rate (FDR) correction was applied to control for multiple comparisons.

### Masks

In our analyses, we used the neuroanatomical guidelines of the Automated Anatomical Labelling Atlas^64^ to define masks for the RE, amygdala, and hippocampus. The masks for functionally defined regions were created using Neurosynth^65^ with the keyword ‘conditioning.’ We created spheres of 8 mm on the following identified peak coordinates: vmPFC (MNIxyz = −2, 46, −10), sgACC (MNIxyz = 0, 26, −12), and dACC (MNIxyz = 0, 14, 28).

### Animal subjects

Adult male and female Long-Evans Blue Spruce rats (200-224 g) purchased from Envigo were used for the experiments. They were acclimated to the vivarium for one day upon arrival and handled 1 min/day for five days before the experiments. They were kept on a 14/10 h light/dark cycle with *ad libitum* access to food and water. All experiments were conducted during the light cycle and all procedures were approved by the Texas A&M University Animal Care and Use Committee.

### Viruses

The AAV8-CaMKII-GCaMP6f virus purchased from Addgene was used for the fiber photometry experiment. The virus was diluted to a final titer of 5 x 10^12^ GC/mL using a Dulbecco’s phosphate-buffered saline solution.

### Surgical procedures

Animals were anesthetized with isoflurane (5% for induction, ∼2% for maintenance). They received 5 mg/kg Rimadyl (i.p.) and were placed into a stereotaxic frame (Kopf Instruments) after the scalp was shaved. Lidocaine HCI (2%) with epinephrine (Cook-Waite) was injected in the scalp (intra-dermal) and povidone-iodine was applied to the skin. An incision was made, the scalp was retracted, and the skull was leveled using the bregma and lambda coordinates.

For the fiber photometry experiment, animals received the virus injection at the midline. A hole was drilled above the RE (AP:-2.1 mm, ML: 1.25 mm, DV:-7.09 mm relative to the bregma surface with a 10° angle from the midline). The virus (0.5 μL) was infused at a rate of 0.1 μL/min using an injector connected to a Hamilton syringe mounted in an infusion pump (Kd Scientific) with polyethylene tubing. The injector tip was kept in place for an additional 10 min and then was retracted. The incision was cleaned, sutured, and a topical antibiotic (Triple Antibiotic Plus; G&W Laboratories) was applied. Approximately 3 weeks later, the rats underwent another surgery for fiber optic placement. Four small holes were drilled in the skull to affix four jeweler’s screws and the hole above the RE was reopened. An optic fiber (10 mm, 200 μM core, 0.39 NA; RWD, China) was implanted using the coordinates AP:-2.1 mm, ML: 1.25 mm, DV:-6.79 mm (0.3 mm above viral injections) relative to the bregma surface with a 10° angle from the midline. The fiber was affixed to the skull with black dental cement (Contemporary Ortho-Jet Powder, Lang Dental).

For the electrophysiology experiment, four small holes were drilled in the skull to affix four jeweler’s screws. One of these screws functioned as a ground for electrophysiological recordings and was positioned posterior to lambda. Another hole was drilled above the RE to insert an electrode array (16-channel; 50 μm, wires spaced 200 μm in a 4 x 4 arrangement; AP:-2.1 mm, ML: 1.25 mm, DV:-7.09 mm relative to the bregma surface with a 10° angle from the midline). After slowly lowering the electrode array, a ground wire was wrapped around the ground screw and coated with a conductive paint to ensure electrical continuity (Silver Print II; GC electronics). The array was affixed to the skull with dental cement.

After surgery, antibiotic ointment (Triple Antibiotic Plus Ointment, Cosette Pharmaceuticals) was applied to the incision. All animals were then returned to the vivarium and allowed to recover for 7-10 days before beginning any behavioral procedure.

### Behavioral procedures

For fiber photometry and electrophysiology recording experiments, animals underwent habituation, auditory fear conditioning, extinction, and extinction retrieval test in the same chambers (30 x 24 x 21 cm, Med Associates, St. Albans, VT). Percentage of time freezing was used as the measure of fear. For the fiber photometry experiment, freezing levels were captured by a video camera recording system and obtained with the Video Freeze software (Med Associates). For the electrophysiology experiment, freezing levels were captured by load-cell displacement in the chamber floors that was directly recorded by the electrophysiological recording software (OmniPlex, Plexon, Dallas, TX).

Distinct contexts (Contexts A, B, and C) were used for habituation, conditioning, and extinction and retrieval. These contexts were created by using distinct transport boxes, bedding, room lighting, chamber floor, odor, and chamber background. For fiber photometric recordings, Context A involved transporting animals with white transport boxes, red room lights, fans, cage lights, and plastic floors placed over the grid floors. In addition, chambers were wiped with 1.0% ammonium solution to provide a distinct olfactory cue. Context B involved transporting animals with black boxes and red room lights. Cage lights and fans were turned off and there were no plastic floors covering the grids. 3.0% acetic acid was used to wipe the chambers. Context C had white transport boxes, white room lights, cage lights, fans, and plastic floors. 70% ethanol was used to wipe the chambers. For electrophysiological recordings, Context A involved white transport boxes with bedding, white room lights with no cage lights, plastic floors, and a black and white circle-shaped background paper taped onto the back wall of the chamber. The cabinet door was half open and 70% ethanol was used to wipe the chamber. Context B had white transport boxes without bedding, red room lights with no cage lights, grid floors, and a background paper containing black stars on white background. The cabinet door was kept closed and 1.0% ammonium was used to wipe the chamber. Context C had black transport boxes with bedding, white room lights with no cage lights, plastic floors, and a black and white striped background. The cabinet door was half open and 3.0% acetic acid was used to wipe the chamber.

Habituation took place in Context A with five tone (CS: 10 s, 80 dB, 2 kHz) presentations separated by 60 s inter-stimulus intervals (ISIs). Conditioning took place in Context B with 5 CS and foot shock (US: 1 s, 1.0 mA) pairings separated by 60 s ISIs. To extinguish fear associated with the conditioning context, rats were exposed to Context B for 20 min across two consecutive days (context extinction and context extinction retrieval), without CS or US presentations. After context extinction, all rats underwent cued extinction in Context C with 45 CS-alone presentations separated by 30 s ISIs. Once extinction memory was acquired, an extinction retrieval test took place in Context C with five CS-alone presentations separated by 30 s ISIs. A 180 s stimulus-free period was provided prior to CS presentations each day to assess baseline levels of freezing. All behavioral testing was separated by 24 h.

### Fiber photometry recordings

Calcium transients were recorded using a fiber photometry system (FP3200, Neurophotometrics Ltd., San Diego, CA). 470 and 410 nm LEDs were used to detect calcium-dependent and calcium-independent (isosbestic) activity, respectively. Light intensities were adjusted to obtain ∼ 50 μW power at the fiber tips. 470 and 410 nm LEDs were interleaved, and 40 Hz sampling rate was used for both 470 and 410 nm LEDs.

### Electrophysiology recordings

Extracellular single-unit activity was recorded using a multichannel electrophysiology system (Plexon, Dallas, TX). Briefly, we recorded wideband signals from each channel. The signals were then amplified (2000x), digitized (40-kHz sampling rate), and saved for offline sorting and analyses. 16-channel (Innovative Neurophysiology, Inc., Durham, NC; 4 x 4 50μm diameter tungsten electrodes, 10.5 mm electrode length, 200 μm electrode spacing, and 200 μm row space) (*n* = 21, 11 males) and 32-channel (Buzsaki32L, NeuroNexus Technologies, Inc.; 50 μm diameter silicon probes, 10 mm length, four shanks with 600 μm spacing with eight electrodes on each) (n = 3, 2 males) multielectrode arrays were used.

### Histology

Animals were overdosed with sodium pentobarbital (Fatal Plus, 100 mg/kg, 0.5 ml, i.p). Electrophysiology experiment animals were given DC current lesions (0.1 mA pulse, 10s) (World Precision Instruments) to mark electrode tips at the corners of the arrays. All animals were transcardially perfused with ice-cold physiological saline and 10% formalin solutions. Brains were obtained and kept in 10% formalin solution for an additional 24h at 4° C. They were then transferred to a 30% sucrose solution until they sank for approximately 3 days. Brains were sliced at-20° C using a cryostat (Leica Microsystems). For the fiber photometry experiment, 30μ-thick sections were mounted and coverslipped using the Fluoromount-G, with DAPI (Invitrogen) mounting medium. Viral expression and fiber optic placements were verified using a Zeiss microscope (Axio Imager).

For the electrophysiology experiment, 30μ-thick sections were mounted on gelatine-subbed slides, stained with thionin (0.25%), and coverslipped using the Permount (Fisher Scientific) mounting medium. Electrode placements were verified using a brightfield microscope (Leica MZFLIII). Animals with missed injection, electrode and fiber placements were excluded from further analyses.

### Data analyses

For the fiber photometry signal, change in fluorescence (ΔF/F) was used as a measure of calcium activity and ΔF/F values were obtained from the raw fiber photometry signal using the open-source analysis software (pMAT)^66^. For electrophysiological data, waveforms exceeding three standard deviations below baseline noise were selected for unit sorting (Offline Sorter, Plexon). Spike sorting was made manually, using 2D principal component analysis after high-pass filtering (600 Hz). Only well-isolated units were included in the analyses. In the case of two units showing similar waveforms and timestamps, only one was included. Sorted waveforms and timestamps were exported to NeuroExplorer (Nex Technologies, Madison, AL) for further analyses. Unless otherwise stated, analyses were based on CS-evoked firing for each behavior day. Firing rates from each neuron were binned in 50 ms increments. This period was normalized with *z*-scores to the baseline (CS-free) period and averaged across CS trials (e.g., 5 trials for conditioning). *z*-score values ≥ 3 in 50 ms bins during the first 250 ms following CSs and USs were considered CS-evoked and US-evoked, respectively. To analyze CS response latencies during early extinction, firing rates were binned in 10 ms increments, normalized to the baseline and the earliest bin at which *z*-score values ≥ 3 are observed was averaged for CS-responsive neurons.

### Hierarchical clustering analysis

Initially, the magnitude of individual neuronal activity was normalized using a logarithmic transformation^67^, followed by *z*-score normalization calculated using the mean and standard deviation during the 1-second preCS onset or pre-freezing period. Subsequently, the data were smoothed over time using a Gaussian filter (σ ≈ 3). A hierarchical cluster tree was then generated using Ward’s method^68^, as previously described^69^, and implemented with the Scikit-learn Python library (version 1.6.9). A cutoff threshold of ∼30% of the maximum dataset value was determined using the Elbow Criterion, which evaluates the total within-cluster sum of squares as a function of the number of clusters. Clusters containing fewer than three units and units not falling into a cluster were discarded.

### Statistics

All data were represented as means ± SEM and *p* < 0.05 was considered statistically significant. GraphPad Prism 9.5.0 was used for behavioral data analyses. Repeated-measures ANOVA was conducted, and significant results were followed by Tukey’s multiple comparisons test. Male and female data were collapsed where significant sex differences were not found. Group sizes were determined based on prior work and the litera-ture.

## Data Availability

The data from these experiments are available from the corresponding author upon request.

## Code Availability

Custom codes used for data analyses are available from the corresponding author upon request.

## Acknowledgements

We thank Drs. Justin Moscarello, Rachel Smith, and Ursula Winzer-Serhan for helpful discussions around the direction and interpretation of this work. We also thank Jaden Carmichael, Mehlum Ali, and Mahima varghese for their help with microscopy. This work was supported by the National Institutes of Health (R01MH065961 and R01MH117852 to S.M.).

## AUTHOR CONTRIBUTIONS

MB and MRM analyzed the fMRI data. TT, MT, and SM designed the electrophysiological experiment; MT and SM designed the fiber photometry experiment. SP and MT performed the fiber photometry experiment; TT performed the electrophysiological experiment. MT and TT analyzed the behavioral and electrophysiological data; MT and SP analyzed the fiber photometry data; FM analyzed the electrophysiological and fiber photometry data; TT, MT, and SM wrote the manuscript.

## SUPPLEMENTARY RESULTS

To determine if individual RE neurons show divergent patterns of activity, we additionally performed hierarchical clustering on normalized firing from each unit, aligned to the CS (CS onset or offset). We performed clustering for early and late extinction, separately. This yielded four different clusters for early extinction (Fig. S1A) and six different clusters for late extinction (Fig. S1D). For both early and late extinction clusters, there were three distinct CS-evoked firing patterns: increase, decrease, and offset change (Fig. S1 B&E). We collapsed clusters showing the same firing pattern and performed a two-way repeated measures ANOVA with CS Onset v Offset and CS v Non-CS as factors (matched for both). This revealed significant main effects of CS v Non-CS for early extinction clusters (Fig. S1C, Offset change: *F*_1,33_ = 23.87, *p* <.0001, Post-hoc for Offset: CS vs Post-CS: *p* =.0153; Decrease: *F*_1,19_ = 50.55, *p* <.0001, and Increase: *F*_1,44_ = 90.56, *p* <.0001, Post-hocs for Onset: Pre-CS vs CS, Offset: CS vs Post-CS: *p*s <.0001) and late extinction clusters (Fig. S1F, Offset change main effect of CS v Non-CS: *F*_1,36_ = 399.8, *p* <.0001, Post-hoc for Offset: CS vs Post-CS: *p* <.0001; Decrease CS v Non-CS x Onset v Offset interaction: *F*_1,15_ = 152.6, *p* <.0001, Post-hocs for Onset: Pre-Cs vs CS, Offset: CS vs Post-CS: *p* <.0001; Increase main effect of CS v Non-CS: *F*_1,43_ = 277.8, *p* <.0001, Post-hocs for Onset: Pre-CS vs CS: *p* <.0001, Offset: CS vs Post-CS: *p* =.0001). These results demonstrate that RE neurons respond to CSs similarly during both early and late extinction. The majority of the RE neurons show matched CS onset and offset responses (both decrease or both increase in firing), with a third population of neurons showing no change in firing during CS onset but offset.

---

**Figure S1.**
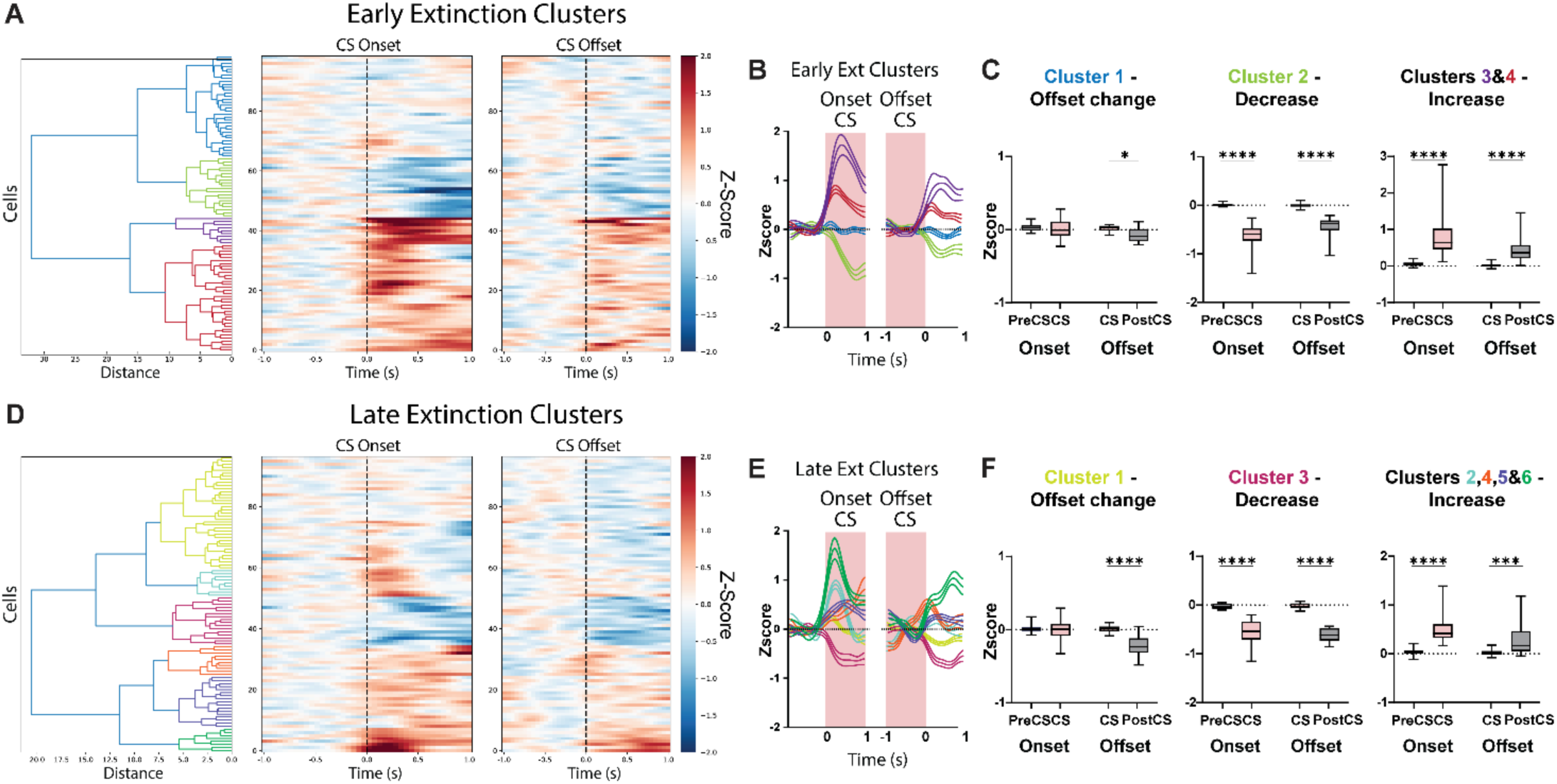
RE neurons respond to CSs similarly during both early and late extinction. Dendrograms showing different clusters based on hierarchical clustering of normalized firing over time during early extinction **(A)** and late extinction **(D)**. Average normalized firing of early extinction **(B)** and late extinction **(E)** clusters over time. **(C,F)** Box and whisker plots showing average normalized firing in CS and non-CS epochs during CS onset and offset. Both early **(C)** and late **(F)** extinction clusters exhibited decreased (middle) or increased (right) firing at CS-onset and-offset (clusters showing the same pattern averaged), while one cluster during both early and late extinction showed no change at the CS-onset but decreased firing at the CS-offset (left, offset change). Data are represented as mean ± SEM. For the context extinction day, we similarly averaged normalized firing from each unit for a-1 s to 1 s period, aligned at behavioral transition (freezing initiation or termination) (see Fig. 6 for context retrieval day). As in the context retrieval day, freezing initiation was associated with a decrease in firing (Fig. S2A, Initiation, Wilcoxon signed rank test, *p* <.0001), with the decrease in firing preceding the freezing initiation. We also observed an increase in firing with freezing termination, although not significant (Fig. S2A, Termination, Wilcoxon signed rank test, *p* =.4437). We then averaged units within animals creating a single value for each animal. A two-way repeated measures ANOVA with Non-freezing v Freezing and Initiation v Termination (matched for both factors) confirmed that firing is significantly lower during Freezing compared to Non-freezing bouts (main effect of Non-freezing v Freezing, Fig. S2B, *F*_1,21_ = 143.3, *p* <.0001; Post-hocs for Initiation: Freezing vs Non-freezing, Termination: Freezing vs Non-freezing, Freezing: Initiation vs Non-freezing: Termination, Freezing: Termination vs Non-freezing: Initiation: *p*s <.0001). We then performed hierarchical clustering on normalized firing from each unit, aligned at behavioral transition (freezing initiation or termination). This again yielded six distinct clusters (Fig. S2C-I). Among these clusters, Cluster 4 and 5 neurons revealed increased firing with freezing as opposed to the dominant pattern (Fig. S2G-H). A two-way repeated measures ANOVA confirmed significantly higher firing for Freezing compared Non-freezing bouts during both freezing initiation and termination (main effects of Non-freezing v Freezing, Fig. S2G right, Cluster 4: *F*_1,14_ = 174.5, *p* <.0001; Fig. S2H right, Cluster 5: *F*_1,14_ = 98.12, *p* <.0001, Post-hocs for Initiation: Freezing vs Non-freezing, Termination: Freezing vs Non-freezing: *p*s <.0001). On the other hand, Cluster 1-3 and 6 neurons revealed decreased firing with freezing (Fig. S2D-F&I), the pattern observed with average normalized firing of all units (Fig. 6A) and fiber photometric recordings (Fig. 3). A similar two-way repeated measures ANOVA confirmed that firing is significantly lower during Freezing compared to Non-freezing bouts for all clusters (main effects of Non-freezing v Freezing, Fig. S2D right, Cluster 1: *F*_1,33_ = 528.9, *p* <.0001; Fig. S2F right, Cluster 3: *F*_1,33_ = 271.6, *p* <.0001; Fig. S2I right, Cluster 6: *F*_1,9_ = 212.7, *p* <.0001, Post-hocs for Initiation: Freezing vs Non-freezing, Termination: Freezing vs Non-freezing: *p*s <.0001; Fig. S2E right, Cluster 2: *F*_1,6_ = 94.85, *p* <.0001, Post-hocs for Initiation: Freezing vs Non-freezing: *p* =.0002, Termination: Freezing vs Non-freezing: *p* =.0089).

**Figure S2.**
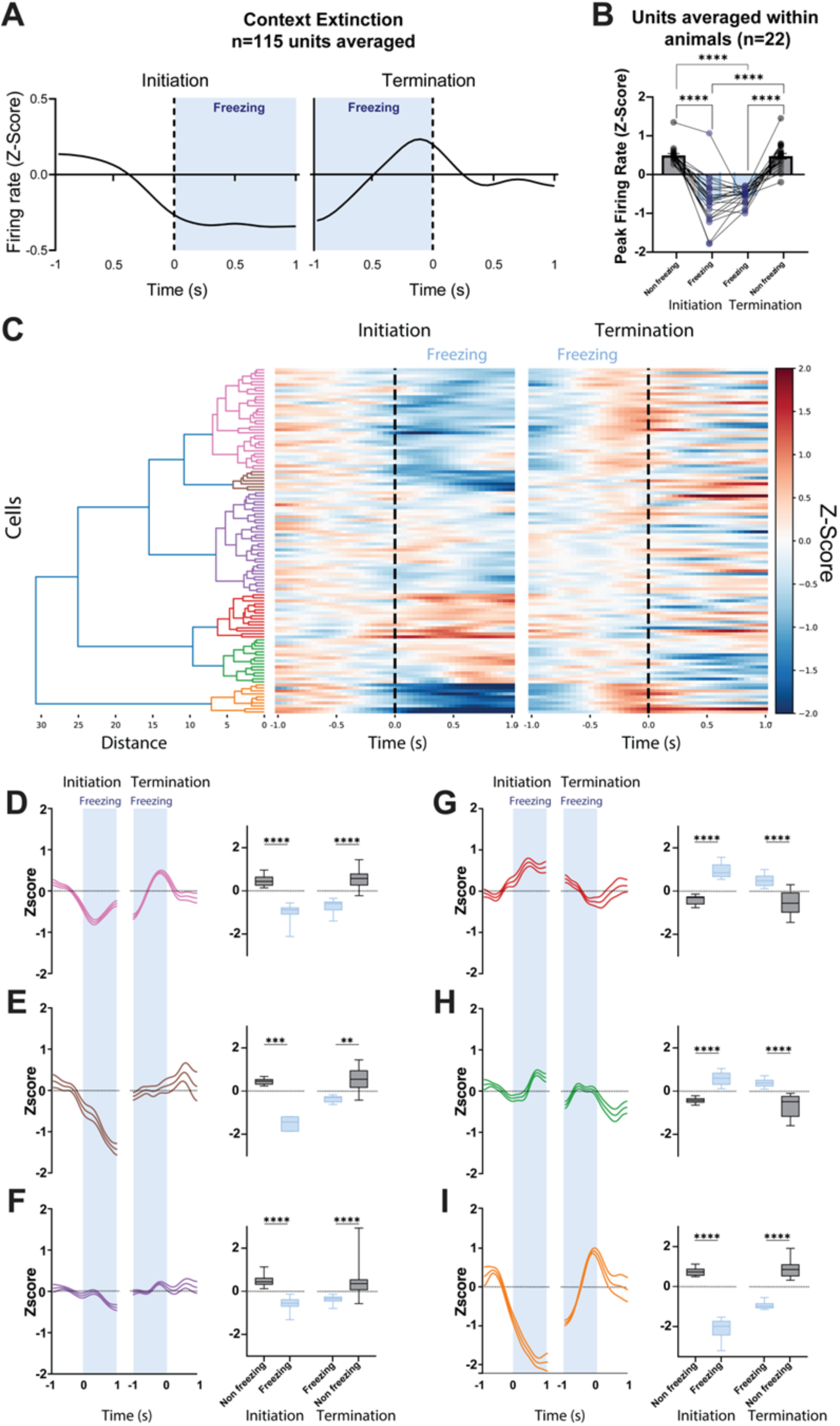
Spontaneous firing of RE neurons tracks behavioral state transitions. **(A)** Average normalized firing of neurons (n = 115) over time during freezing initiation (left) and termination (right). **(B)** Bar graph with peak normalized firing of neurons in non-freezing and freezing epochs during freezing initiation and termination. **(C)** Dendrogram showing six different clusters based on hierarchical clustering of normalized firing over time. **(D-I)** Average normalized firing of each cluster over time (left) and box and whisker plots (right) showing peak normalized firing in non-freezing and freezing epochs during freezing initiation and termination. Data are represented as mean ± SEM.

